# Inception in visual cortex: *in vivo-silico* loops reveal most exciting images

**DOI:** 10.1101/506956

**Authors:** Edgar Y. Walke, Fabian H. Sinz, Emmanouil Froudarakis, Paul G. Fahey, Taliah Muhammad, Alexander S. Ecker, Erick Cobos, Jacob Reimer, Xaq Pitkow, Andreas S. Tolias

## Abstract

Much of our knowledge about sensory processing in the brain is based on quasi-linear models and the stimuli that optimally drive them. However, sensory information processing is nonlinear, even in primary sensory areas, and optimizing sensory input is difficult due to the high-dimensional input space. We developed *inception* loops, a closed-loop experimental paradigm that combines *in vivo* recordings with *in silico* nonlinear response modeling to identify the Most Exciting Images (MEIs) for neurons in mouse V1. When presented back to the brain, MEIs indeed drove their target cells significantly better than the best stimuli identified by linear models. The MEIs exhibited complex spatial features that deviated from the textbook ideal of V1 as a bank of Gabor filters. Inception loops represent a widely applicable new approach to dissect the neural mechanisms of sensation.

---

Since the work of Adrian and Hartline (*1, 2*), finding stimuli that optimally drive neurons has been fundamental for understanding information processing in the brain. In linear systems, the most effective stimuli for eliciting responses match the linear filters. For instance, quasilinear models with center-surround filters have high predictive power in the retina (*3*), and these patterns also strongly drive retinal activity. In primary visual cortex (V1) the most effective stimuli — obtained by *assuming* a linear model — resemble Gabor patterns (*4*). However, the predictive power of the corresponding quasi-linear models is low (*5*), especially when predicting responses to natural stimuli in mice (*6, 7*). This casts doubt on whether Gabor patches are really the optimal stimuli for neurons in mouse V1, and therefore on whether the Gabor filter bank model is the right framework for understanding V1 computations.

In general, identifying the optimal sensory input is inherently difficult due to the nonlinear sensitivity of neurons and the intractably high-dimensional space of possible images. Proposed active learning approaches (*8–11*) are impractical, because experimental constraints limit the number of responses that can be measured from any cell. However, recent progress in deep learning has yielded stronger predictive nonlinear models of neuronal responses (*6, 7, 12–14*). Here we use these models for improved stimulus design *in silico*.

We designed a closed-loop experiment that combines *in vivo* recordings with *in silico* modeling to perform ‘inception’ (*15*)—implanting a particular response in the brain (Fig. 1). Briefly, on day one of an inception loop experiment, we recorded the neural responses of large neuronal populations to a large set of natural images; trained a convolutional neural network (CNN) to predict these responses based on the presented images (*7, 12–14*); and optimized images to maximize the model responses (*16*). Over subsequent days, we presented these tailored images to the corresponding real neurons in the brain to test whether they indeed produce the strongest responses amongst all control stimuli.

**Figure 1:**
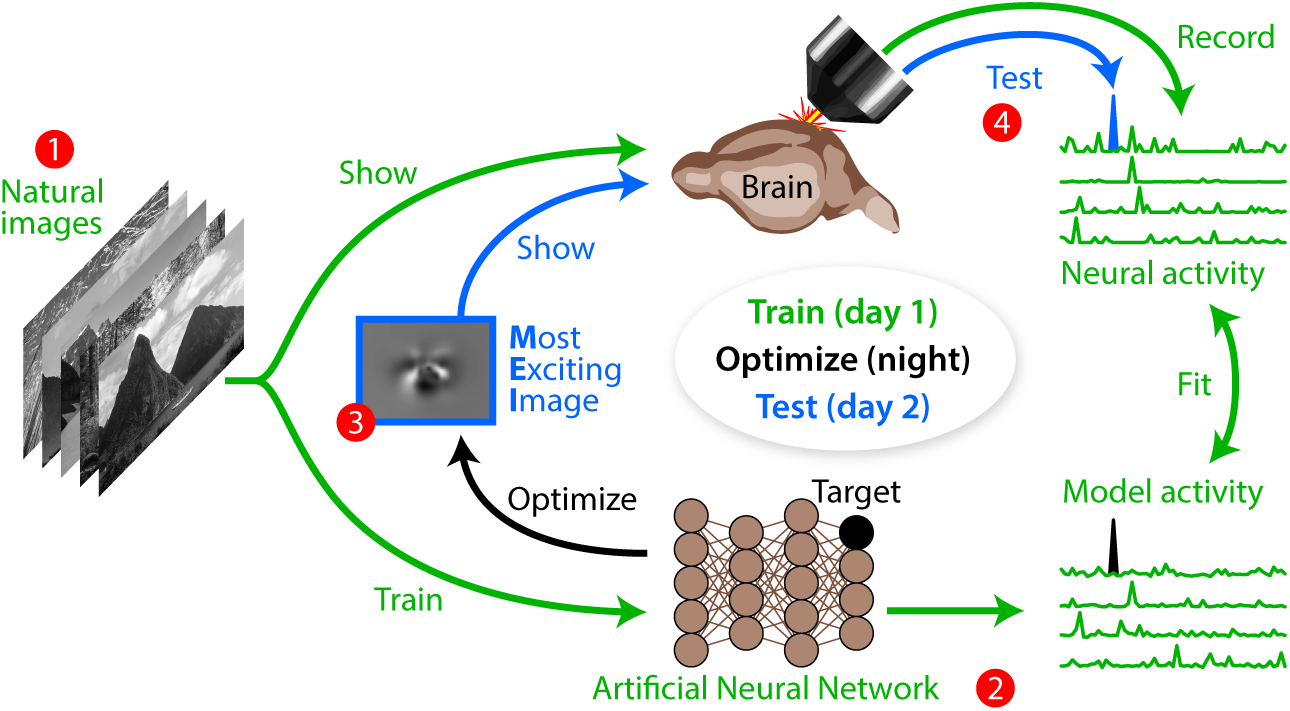
Schematic of the inception loop. On day 1 (green) we showed sequences of natural images to a mouse and recorded neural activity by two-photon calcium imaging. We trained artificial neural networks to reproduce those measured neural responses. Overnight (black) we generated Most Exciting Images (MEIs) for each target neuron in the neural network, as well as for a linear model. On day 2 (blue) we showed these MEIs from nonlinear and linear models back to the same neurons in the brain and compared the responses.

We recorded the responses of more than 2100 neurons in layer 2/3 of primary visual cortex (V1 L2/3) to natural images in two awake mice using two-photon imaging with a large-field of-view mesoscope (Fig. 2A) (*17*). Before each functional imaging session, we recorded a high-resolution anatomical 3D stack (Fig. 2B). Later we registered all recording planes into each of these stacks to identify the neurons of interest across multiple days ((Fig. 2C); see Supplementary Methods for details).

**Figure 2:**
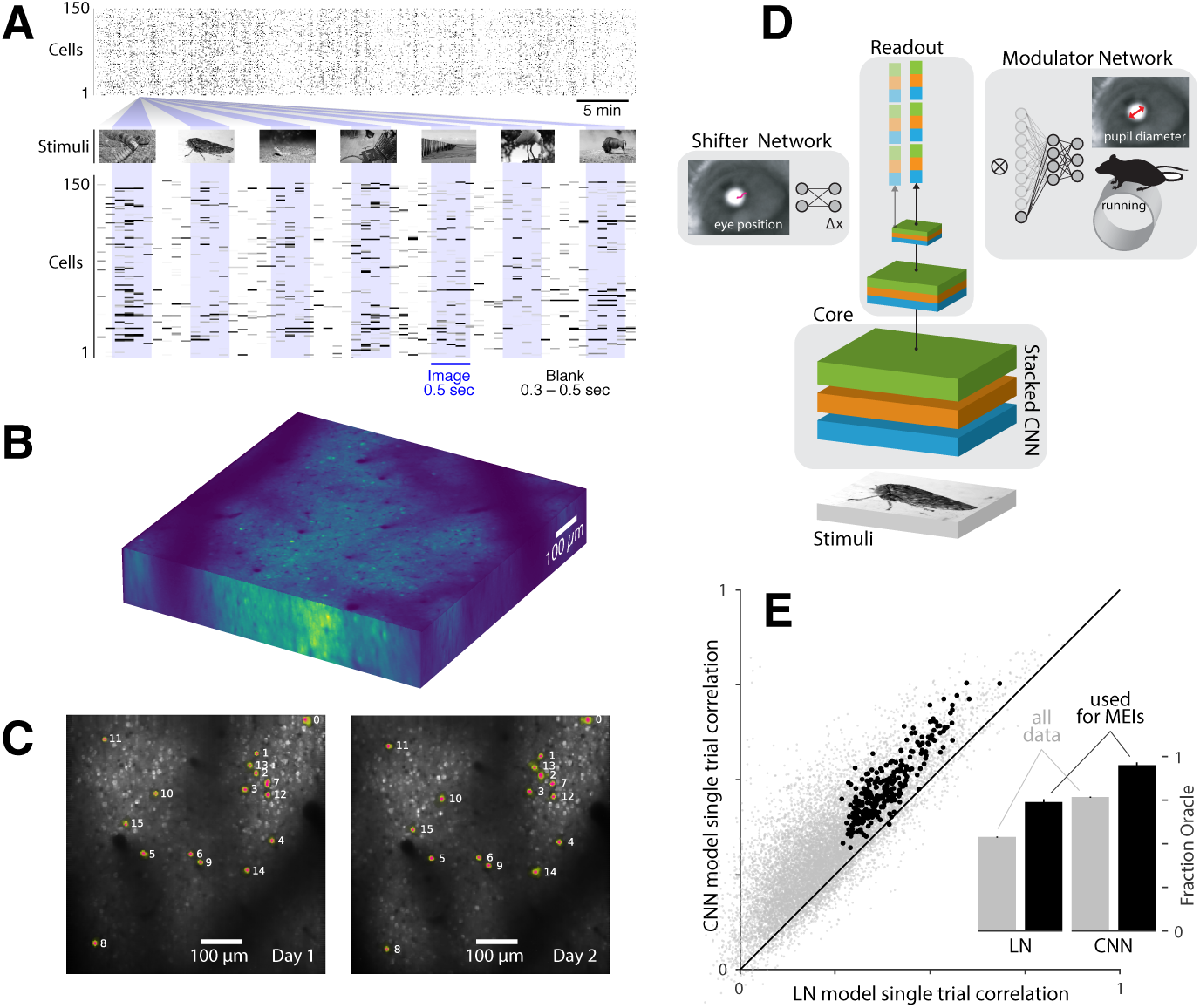
Experimental paradigm and model. **A**: 5100 unique natural images were presented to an awake mouse for 500 ms interleaved with gray screen gaps of random length between 300 – 500 ms. A subset of 100 images were repeated 10 times each to estimate the reliability of neuronal responses (see Methods). Neuronal activity was recorded at 8 Hz in V1 L2/3 using a wide field two-photon microscope. **B**: To identify the same cells across days, structural stacks were recorded on each day and aligned between days. **C**: Cells were reliably matched between days (left *versus* right) by aligning the recording planes into each stack. The two panels show example recording planes on separate days with a subset of the cells used to generate MEIs (colored masks). Cells with identical numbers were matched. **D**: Convolutional neuronal network (CNN) trained to predict neuronal responses: Each network consisted of a convolutional core computing nonlinear features from the image, a readout predicting the neuronal responses from the core, a shifter network accounting for eye movements, and a modulator network predicting an adaptive gain for each neuron based on behavioral variables (see Methods). **E**: CNN *versus* linear-nonlinear (LN) model performance. Each point denotes the correlation between the model predictions and single trial responses. The CNN model significantly outperforms the LN model (paired *t*-test, *t*(12956) = 127.43 with *p ≪* 10^-4^). Black points depict the performance for neurons used to generate MEIs, while grey points designate remaining neurons. Data from both mice are combined in this plot. Inset: Performances for models with frozen shifter and modulator signals, relative to an upper bound estimated from repeated presentations of identical stimuli (oracle).

On the first day we collected the population responses to a set of 5000 unique natural images, which were used to fit a predictive model of neural responses to visual stimuli (Fig. 2A). Another set of 100 images, repeated 10 times each, was used as the test set for the model and for evaluating response reliability. We focused purely on spatial characteristics of the neurons and ignored temporal dynamics. Each image was shown for 500 ms and we extracted each neuron’s response by integrating the deconvolved fluorescence trace over a time window of 50 ms to 550 ms after image onset.

Next, we trained a deep convolutional neural network (CNN) to predict the recorded responses (Fig. 2D). Recent work has established that deep CNNs outperform classical models of V1 such as linear-nonlinear, subunit or energy models (*14, 18–20*). We used a core network that consisted of three convolutional layers shared among all recorded neurons, followed by a neuron-specific linear readout stage. The network also accounted for eye position and behavioral state (running, pupil dilation) of the animal (*7*), which we could measure but not control experimentally. In line with earlier work, we found that the CNN model outperformed a linear-nonlinear model, achieving 77% of the theoretically achievable performance given the noise ceiling in the recordings (Fig. 2E, pooled over models for both mice).

Our final step for day 1 was to obtain the Most Exciting Images (MEIs) for a subset of neurons whose responses were reliable and reasonably well-predicted by both the CNN and the linear-nonlinear (LN) model. We achieved this via a simple optimization procedure (*16*): To find the image that maximally excites a target neuron, we started with a random image and performed regularized gradient ascent until convergence (Fig. 3A). We repeated the optimization for an LN model to get a corresponding estimate of the linear receptive field (RF). After the optimization procedure, we matched the average luminance and contrast between MEI and RF to ensure a fair comparison.

**Figure 3:**
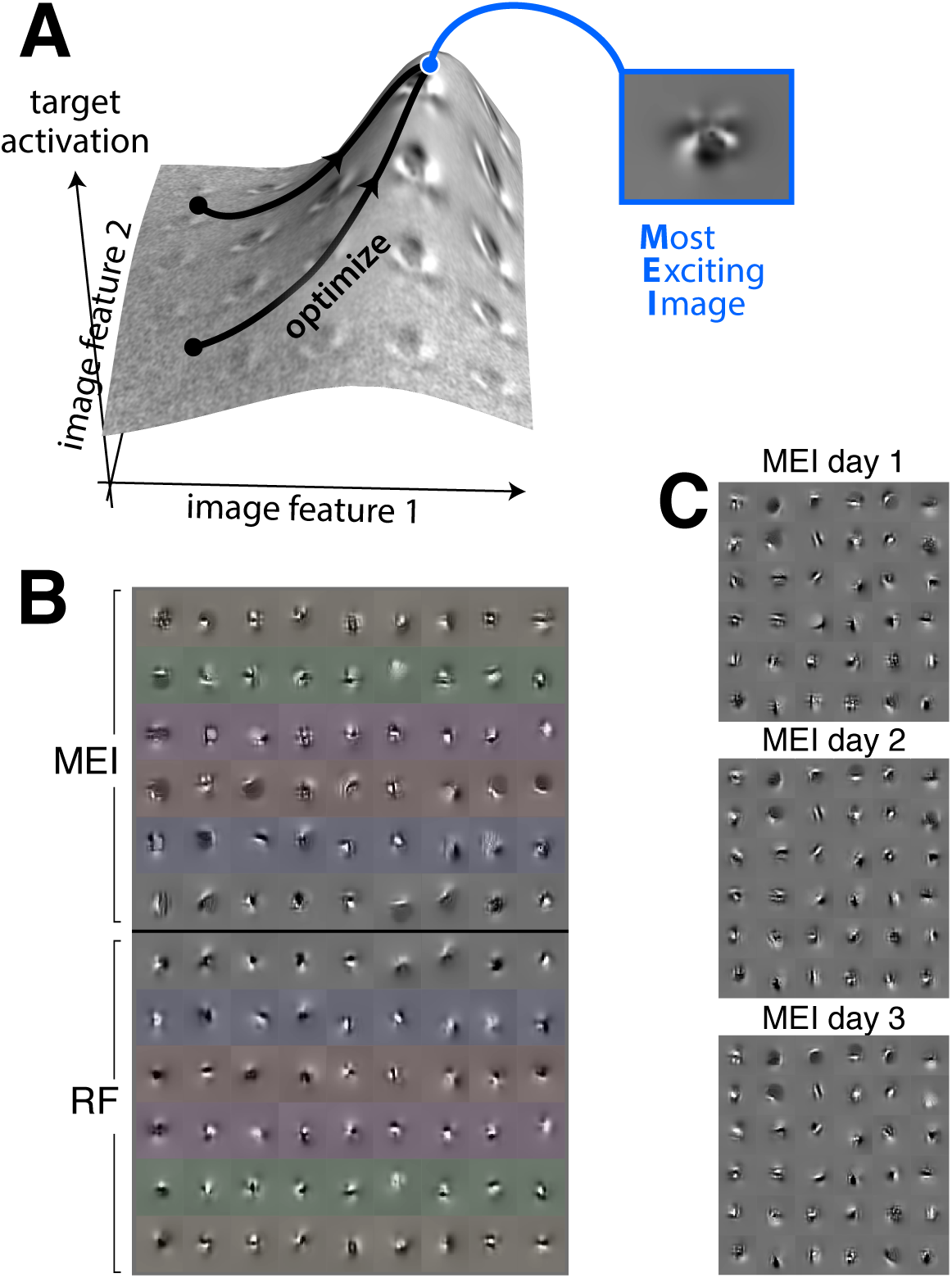
Most Exciting Images (MEI). **A**: Illustration of optimization over all possible images. Vertical axis represents activation of a model neuron as a function of two example image dimensions. Black curves depict optimization trajectories converging to the same MEI from different initializations. **B**: Blocks of example MEIs and linear RFs. The two blocks are symmetrically arranged and tinted to allow for easier visual comparison between matching pairs. MEIs exhibit more complex, higher frequency features than linear RFs. **C**: MEIs are stable across days. Each block shows the MEIs of matched cells computed from models trained to predict natural image responses from scans on three separate days.

The resulting MEIs deviated substantially from the textbook notion of Gabor-shaped V1 receptive fields. They exhibited complex spatial features such as sharp corners, checkerboard patterns, irregular pointillist textures, and a variety of curved strokes (Fig. 3B, Fig. S5). They were often slightly larger than the RFs for the same neurons, and included notably more high frequency details (Fig. S2) that were stable across days (Fig. 3C). While some RFs also exhibited atypical structure, consistent with the conventional wisdom many looked qualitatively more similar to Gabor filters (Fig. 3B, Fig. S4).

We verified that the intricate differences between MEIs and linear RFs reflect actual selectivity of the neurons rather than an artifact of the model (*i. e.* overfitting) by performing a follow-up experiment in the same animal on the same neurons. This time, we showed both MEIs and linear RFs for a subset of 150 neurons from the same neuronal population that had been recorded in the original session (see Supplementary Methods). This experiment confirmed the model’s predictions about the neurons’ responses to MEIs, which were images that had never been seen during training. First, MEIs were highly specific to their target neurons: they evoked consistently higher activity than other neurons’ MEIs (Fig. 4A and Fig. S6, left). The same was true for RF stimuli (Fig. 4A and and Fig. S6, right). Second, responses to MEIs were generally very sparse, with only a few images activating neurons above baseline for both model and measured neurons (Fig. 4B). Third, the model’s predictions correlated strongly with the observed responses to MEIs (Fig. 4C; average correlation coefficient: 0.76).

**Figure 4:**
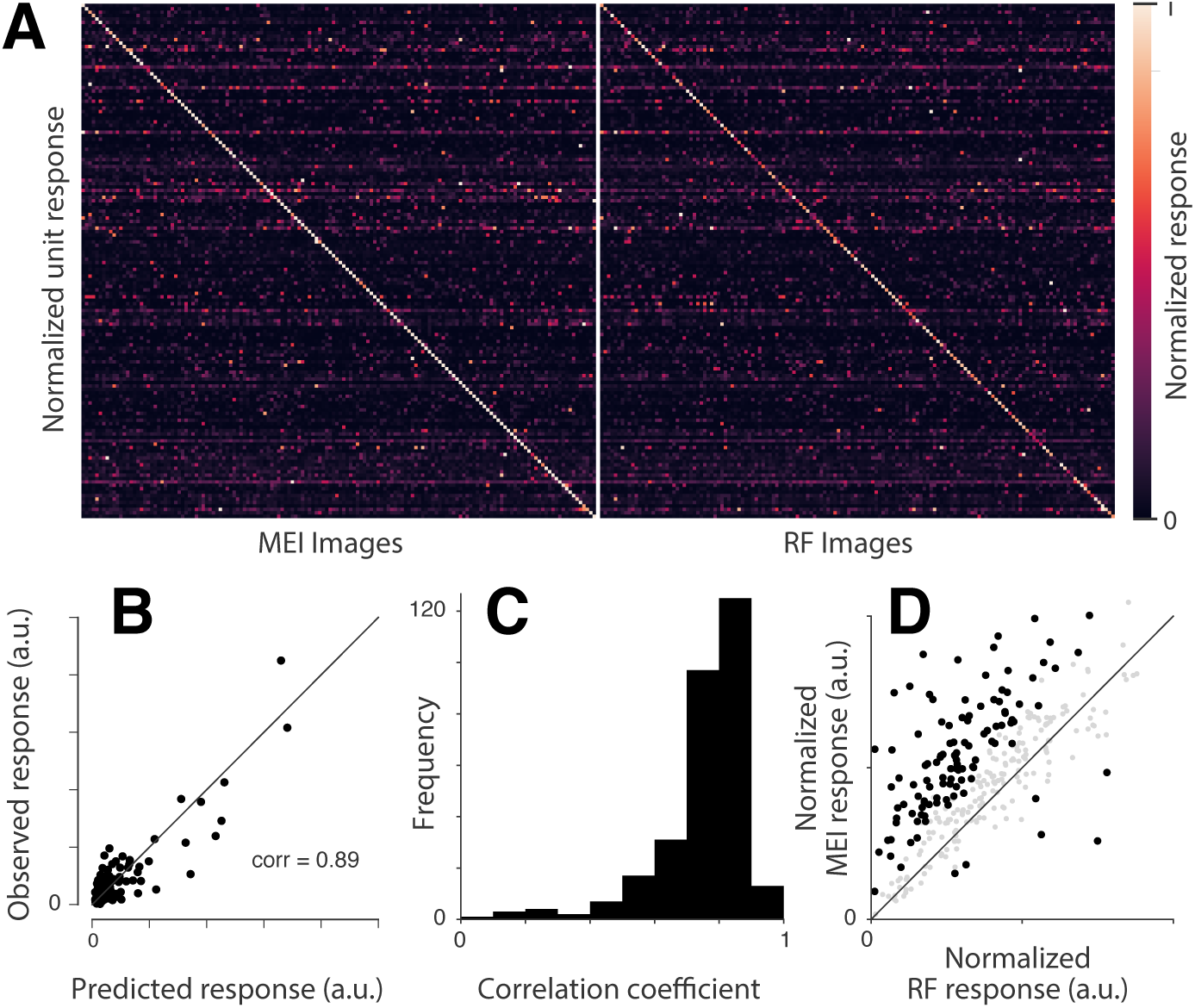
MEIs reliably drive neurons more than linear RFs. **A**: MEIs and linear RFs activate neurons with high specificity: Confusion matrices show the responses of each neuron to the MEIs (left) and RFs (right) of all neurons. Each row in both plots are normalized to the maximum response of the neuron across all presented images. **B**: Predicted *versus* observed responses to all MEIs for one example neuron. Each neuron’s responses to all images were normalized to a single common scale and then pooled across days. **C**: Single trial correlations between the model predictions and the neuronal responses to all presented MEIs for all neurons used for MEI generation. **D**: MEIs activate neurons more strongly than RFs. Each point corresponds to the activity of a single neuron in response to its RF or MEI, averaged across 38–60 repeated presentations. Neurons with significant difference in their mean responses are colored black. 235 out of 300 neurons showed a stronger response to their MEIs than their own RFs, of which 107 showed a statistically significant effect (*p <* 0.05, two tailed Welch’s *t*-test). In contrast, only 6 had a statistically significantly stronger response to RFs, consistent with random chance. Overall MEIs activated their target neurons significantly stronger than the corresponding RFs (Wilcoxon Signed-Rank test, *Z* = 5816.0, *p ≪* 10^-4^ over *n* = 300 neurons pooled across both mice).

Most importantly, MEIs evoked significantly stronger activity than the linear RFs of the corresponding neurons (Fig. 4D). The striking specificity and repeatability of individual neuronal responses to the predicted stimuli validate the CNN model and demonstrate that the MEIs reflect the true selectivity of the neurons better than their linear RFs.

Comparing MEIs to linear RFs is a stringent test of current standard models of V1, including nonlinear extensions such as energy models (*21*). For instance, the optimal stimulus for an energy model is any linear combination of the two filters in the quadrature pair (*i. e.* a Gabor); based on these models, the linear RF and MEI should be identical.

Our work shows that these high-performing CNN black-box models of the visual system do generalize and can make *in silico* inferences about non-trivial computational properties of V1 neurons. We find that even V1 neurons prefer features that are more complex than the classic orientated edges described by Hubel and Wiesel (*22*) and predicted by many theories of early visual processing (*23*). These complex feature selectivities in V1 could provide a faster, if less general, way to extract task-relevant causal variables from natural inputs. This might be beneficial for small animals like mice, whose diminutive size may impose stringent constraints on computation.

Strong predictive models allow for a nearly unlimited number of *in silico* experiments that can be tailored to individual neurons or populations. This could be especially important when studying high-dimensional tuning properties, such as invariances (*18, 24*) or equivariances (*25*): searching directly for these dimensions in the brain is slow and costly, and is unlikely to reveal the relevant stimulus manifolds. Naturally, predictions of these models require experimental verification. Our current work with inception loops shows that such verification is indeed feasible.

Inception loops provide many opportunities for future neuroscience studies. We performed this experiment in V1 because it constitutes a particularly stringent test, due to the large existing literature on V1 receptive field structure. In extrastriate or non-visual areas which we understand much less well, this approach (*26–28*) might prove even more useful for revealing feature selectivities. Finally, this method could be used not only to target the activity of single neurons, but entire populations of neurons; not only to maximize responses, but to control spatiotemporal patterns; and not just to optimize using inputs at the sensory periphery, but through any type of causal manipulation including patterned optogenetic stimulation. Combining such emerging neurotechnologies with modeling and machine learning in a closed inception loop would provide a level of control that would be a far more powerful tool to probe and evaluate brain transformations than we currently have. This will likely lead to a much richer understanding of neural computation, as well as to improved interventions for clinical applications.

## Acknowledgments

We thank George Denfield for comments on the manuscript. Supported by the Intelligence Advanced Research Projects Activity (IARPA) via Department of Interior/Interior Business Center (DoI/IBC) contract number D16PC00003. The U.S. Government is authorized to reproduce and distribute reprints for Governmental purposes notwithstanding any copyright annotation thereon. Disclaimer: The views and conclusions contained herein are those of the authors and should not be interpreted as necessarily representing the official policies or endorsements, either expressed or implied, of IARPA, DoI/IBC, or the U.S. Government. Fabian Sinz is supported by the Institutional Strategy of the University of Tübingen (Deutsche Forschungsgemeinschaft, ZUK 63) and the Carl-Zeiss-Stiftung.

## Supplementary materials

### Materials and Methods

#### Neurophysiological experiments

All procedures were approved by the Institutional Animal Care and Use Committee (IACUC) of Baylor College of Medicine. Briefly, two male mice aged 67 and 70 days and expressing GCaMP6s in excitatory neurons via SLC17a7-Cre and Ai162 transgenic lines (Jackson IDs 023527 and 031562, respectively) were anesthetized and a 4 mm craniotomy was made over visual cortex as previously described (Reimer et al., 2014, Froudarakis et al., 2014). Mice were head-mounted above a cylindrical treadmill and calcium imaging was performed using Chameleon Ti-Sapphire laser (Coherent) tuned to 920 nm and a large field of view mesoscope (*17*) equipped with a custom objective (0.6 NA, 21mm focal length). The laser power after the objective was always kept below 60 mW. Visual stimuli were presented to the left eye with a 15” TFT monitor and resolution 2560 *×* 1440 px positioned 15 cm from the eye. Rostro-caudal treadmill movement was measured using a rotary optical encoder with a resolution of 8000 pulses per revolution. We used the light diffusing from the laser during scanning through the pupil of the animal to capture eye movements. The images of the pupil were reflected through a hot mirror and captured with GigE CMOS camera (Genie Nano C1920M, Teledyne Dalsa) at 20 fps and 1920 1200 px resolution. The contour of the pupil was extracted semi-automatically for each frame, and the center and the major radius of a fitted ellipse were used as the position and the dilation of the pupil.

Pixelwise response across a 2400 *×* 2400 *µ*m region of interest (0.2 px/*µ*m) at 200 *µ*m depth from the cortical surface to drifting bar stimuli was used to generate a sign map for delineating visual areas (*29*). An imaging site was chosen in primary visual cortex with minimal blood vessel occlusion and maximal stability. The craniotomy window was leveled with regards to the objective with six degrees of freedom, five of which were locked between days to allow return to the same imaging site using the z axis. Imaging was performed at 8 Hz for all scans, using a remote objective to sequentially image ten 630 *×* 630 *µ*m fields per frame at 0.4 px/*µ*m xy resolution. Fields were spaced 5 *µ*m apart in depth to achieve dense imaging coverage of a 630 *×* 630 *×* 50 *µ*m^3^ xyz volume, with the most superficial plane positioned around 200 *µ*m from the surface in layer 2/3. Cells in the imaged volume were oversampled at this z resolution, often corresponding to three or more imaging planes per cell and allowing matching across days with *≤* 2.5 *µ*m vertical distance between masks. Imaging data were motion corrected, automatically segmented and deconvolved using the CNMF algorithm (*30*), then further selected by a classifier trained to detect somata based on the segmented cell masks but not according to visual responsiveness. This resulted in 5300 to 8400 soma masks per scan. A structural stack encompassing the volume and imaged at 0.6 *×* 0.6 *×* 1 px/*µ*m xyz resolution with 100 repeats was used to register the scan average image into a shared xyz frame of reference between scans (see below).

#### Monitor positioning across days

In order to have a consistent monitor placement relative to the mouse across all imaging sessions, we placed the aggregate receptive field for each session on the center of the monitor. To map the receptive field we used ∼ 5º single dark dots over bright background stimuli tiling the center of the screen in a 10 *×* 10 grid and averaged the ∼ 0.5 – 1.5 sec after the stimulus onset of the calcium trace of all central pixels of the imaged volume (∼150*×*150*µ*m) across all repetitions of the stimulus for each location. We fitted the resulting 2D map using an elliptic 2D Gaussian and we centered the position of the receptive field by displacing the monitor with ∼ *±*2.5º precision between imaging sessions.

#### Cell registration across days

We registered each 2D plane to the 3D stack using an affine transformation matrix with 9 degrees of freedom estimated via gradient ascent on the correlation between the average recorded plane and the extracted plane from the stack. We matched cells across scans using their estimated centroids in the stack: matching each cell of interest to the closest cell in the stack from a different scan. For any two scans in a mouse we used multiple stacks and matched pairs that had the highest number of matches across stacks. To functionally confirm that cells were well matched across days we correlated the average responses to the repeated images across days. The resulting confusion matrix is strongly diagonal demonstrating that matched cells across days have the same response properties (Fig. S1).

#### Natural stimuli and data transformation for network training

Stimuli consisted of 5100 natural images from ImageNet (*31*), cropped to fit a 16:9 monitor aspect ratio, and converted to gray scale. In each scan, we showed 5000 unique images and 100 images repeated 10 times each. Each image was presented for 500 ms followed by a blank screen lasting between 300 ms and 500 ms, sampled uniformly from that range.

#### Preprocessing of neural and behavioral data

The neural responses were first deconvolved using non-negative matrix factorization (*30*). We subsequently extracted the accumulated activity of each neuron between 50ms and 550ms after stimulus onset using a Hamming window. Behavioral traces, used as auxiliary signals in model training (see below), were extracted using the same temporal offset and integration window. For training the networks, images were isotropically downsampled by a factor of four to 64 *×* 36 pixels. Input images, the target neuronal activities, and pupil positions were normalized across training set by subtracting the mean and dividing by the standard deviation. All behavioral traces were divided by the standard deviation over the training set.

**Figure S1:**
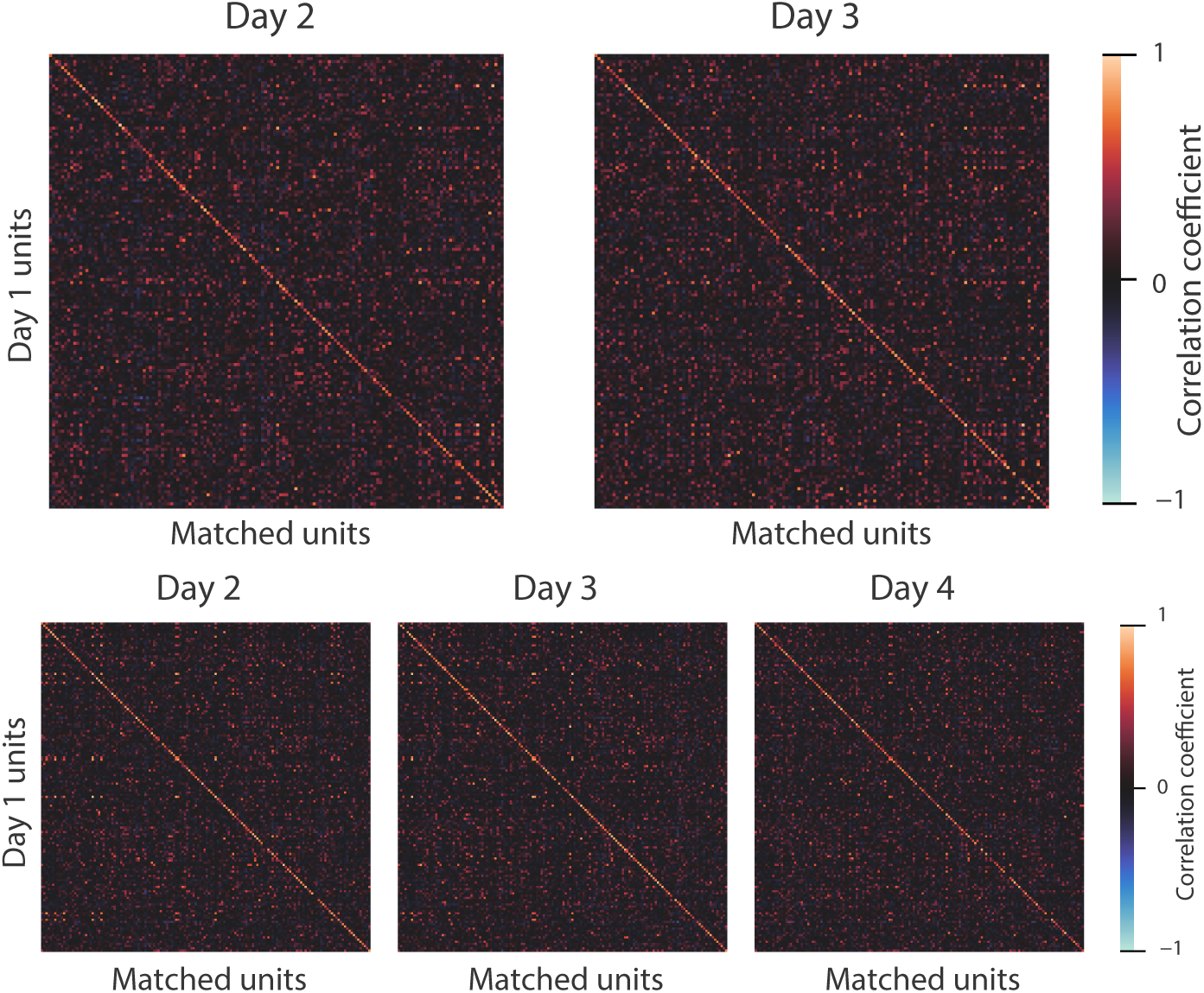
Cross-correlation matrix of the average response to the repeated images across days for matched cells for Mouse 1 (top row) and Mouse 2 (bottom row). The confusion matrices are strongly diagonal, demonstrating that matched cells share the same functional properties.

#### Network Architectures

The network considered here consists of three base elements (Fig. 2D): a common *core* for all neurons providing nonlinear features computed from static images, a dedicated *readout* mapping the core features to the responses for each neuron, a *modulator network* predicting a gain factor for each neuron depending on the running state and the pupil dilation of the animal recorded during the experiment, and an additional *shifter network* predicting receptive field shifts from pupil position changes tracked in an eye camera video (Fig. 2D).

The linear-nonlinear (LN) model and the nonlinear (CNN) model differ only in their core: The CNN core consists of a three layer convolutional network with skip connections. Each layer in that network consists of a convolution layer without bias, a batch norm layer (*32*), and an ELU nonlinearity (*33*). All layers are stacked along the channel dimension before the activations are fed into the readout (see the feature extraction network in (*7*)). The core for the linear-nonlinear (LN) network is identical to the CNN core except that all nonlinearities were removed. This ensures maximal similarity between the two networks in terms of networks components, maximal receptive field size, or behavioral modulation.

For the readout, we model the neural response as an instantaneous affine function of the core followed by an ELU nonlinearity and offset of 1 to make the response positive. For each image, the output of the core is a tensor *v ∈* ℝ^*w×h×c*^. We model the location of each neurons receptive field with a spatial transformer layer using a single grid point reading from *v*_*ijk*_ (*7,34*), i.e. we extract a vector **v**_*x*_*i*_ *y*_*i*_:_ *∈* ℝ^*c*^ via bilinear interpolation of neighboring pixels of the location (*x*_*i*_, *y*_*i*_) in the tensor *v*. To learn the grid points via gradient descent, we decompose *v*_*ijk*_ into several spatial scales through repeated application of an 5 *×* 5 Gaussian lowpass filtering: **v**^(*j*)^ = lowpass^(*j*)^(**v**). Like in a Gaussian pyramid, we keep the difference between the filtered component and the channel. However we included model variants in which, unlike in a Gaussian pyramid, we did not downsample the feature image explicitly since we empirically found that to perform slightly better on some datasets. The exact variant used was determined by model selection on a validation set (see Training and Model Selection below).

The spatial transformer layer then extracts *ℓ* feature vectors from the same relative location (*x*_*i*_, *y*_*i*_) at each scale and stacks them into a single feature vector of dimension *k × ℓ* fed into the final affine function and nonlinearity 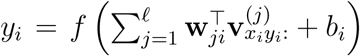 (Figure 2D). The relative spatial location is shared across scales.

To account for fluctuations in neural responses unrelated to the visual stimulus, we use pupil dilations and their temporal derivative, as well as the absolute running speed of the animal which are known correlates of brain state changes (*35–37*). We use these three variables to compute a single gain factor for each neuron that scales the output of the readout layer. We predict the gain using a multi-layer fully-connected perceptron (MLP) with ReLU nonlinearities at all hidden layers and an exponential nonlinearity at the last layer to enforce positivity.

Unlike primates, mice are hard to train to fixate their gaze in a single position, complicating eye tracking. To model the responses of thousands of neurons in a free viewing experiment, we directly estimate a receptive field shift for all neurons from the tracked pupil position solely based on optimizing the predictive performance of the network (*7*). Note that the pupil location is measured in coordinates of the camera recording the eye, while the shift needs to be applied in monitor coordinates. This transformation can either be estimated by a calibration procedure (*38–40*), or learned from the data using regression on pairs of eye camera–monitor coordinates. We use a one layer perceptron (MLP) with a tanh nonlinearity for predicting the joint receptive displacement for all neurons.

#### Training and Model Selection

All experiments and analyses were performed using Data-Joint (*41*), Numpy/SciPy (*42*), Matplotlib (*43*), Seaborn (*44*), Jupyter (*45*), PyTorch (*46*), and Docker (*47*). We trained two different network architectures: a linear-nonlinear (LN) model and a CNN model. Both models used shifter and modulator networks, and a point readout as described above. For any particular configuration of the network, we trained four instances of each network, corresponding to four different random seeds that determined the random initialization. On all but the first convolutional layer, we used a group sparsity regularizer for all weights corresponding to one output channel. The first layer was regularized by penalizing the L2 norm of the 3 *×* 3 Laplace-filtered weights, weighted by an inverse Gaussian profile of the form: 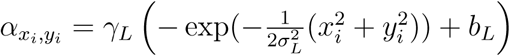, where *x*_*i*_ and *y*_*i*_ are the spatial pixel positions of the convolutional weights relative to the center of the filter. The Gaussian profile encourages the filter to be smoother at the edge. Empirically, we observed this profile also to help center the receptive field within the convolutional kernel’s spatial extent. The values of *σ*_*L*_ and *b*_*L*_ were set to be 0.5 and 1, respectively. All readout weights were penalized by an *L*_1_ sparsity regularizer with weight *γ*_*r*_. The values of *γ*_*L*_ and *γ*_*r*_ were chosen from a discrete set of values as part of the hyperparameter search (Table 1).

**Table 1:** Possible values of hyperparameters during model selection.

| Symbol | Description | Possible Values |
| --- | --- | --- |
| $\gamma_L$ | regularization constant for first layer | $\{50, 100\}$ |
| $\gamma_r$ | L1 regularizer on readout features | $\{0.1, 1.0\}$ |

The dataset was split into training, validation, and test set. The test set consisted of 100 images with 10 repeats each. The remaining unique 5000 images were randomly split into 4477 training images and 523 validation images. The network was trained on the training set using Poisson loss 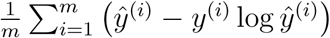 where *m* denotes the number of neurons, *ŷ* the predicted neuronal response, and *y* the experimentally recorded one. We used early stopping on the correlation between predicted and measured neuronal responses on the validation set (*48*). If the correlation did not increase within the last 10 epochs (passes through the entire training set), training was halted and the model was set to the best-performing model over the course of training. After the first stop, training was resumed after we decreased the learning rate from 5 *×* 10^-3^ to 10^-3^, and continued until training was stopped again. The network was trained using the standard stochastic gradient descent technique with the Adam optimizer (*49*) using a batch size of 60. Once the training was completed, the trained network was evaluated on the validation set to yield the final score which served as the basis for hyperparameter selection. The exact configuration of the architecture, including the size of the convolution kernels, the weights of regularizers, or whether to downsample the spatial pyramid, were determined using a grid search model selection on the validation performance.

After the hyperparamter search, the best CNN model for Mouse 1 was obtained with *γ*_*L*_ = 100, *γ*_*r*_ = 0.1, and no downsampling on readout layer. For Mouse 2, the best model had *γ*_*L*_ = 100, *γ*_*r*_ = 0.1, and downsampling on the readout layer. The best LN model for Mouse 1 used *γ*_*L*_ = 50, *γ*_*r*_ = 1.0, and no downsampling on readout layer. For Mouse 2, the best model used *γ*_*L*_ = 100, *γ*_*r*_ = 0.1, and downsampling on the readout layer. Other aspects of the network configurations, namely, the input convolution kernel size *k*_in_ = 15, the number of hidden layer *n* = 2, hidden layer convolution kernel size *k*_hidden_ = 7, and number of channels *c* = 32, were manually selected based on previous experience on other datasets when developing the model architecture.

Since the imaging volume was densely scanned (10 planes spaced 5 *µ*m in depth), cell masks in several planes potentially corresponded to the same soma. We did not take any special measures to merge such overlapping masks during the training but trained the network on all 5335 and 7622 unit activities for Mouse 1 and Mouse 2, respectively. Special care was taken during the unit selection for MEI generation to avoid generating MEIs targeting two or more masks that belong to the same neuron. For the subsequent step of synthetic image generation, we used the best CNN and LN model for each mouse along with their starting seed variants.

#### Oracle correlation and fraction of oracle

Cortical neurons naturally exhibit substantial response variability. To estimate a bound for the maximally achievable correlations, we estimated an “oracle” by correlating the leave-one-out mean across repeated images in the test set with the remaining trial, and averaged that correlation across different images and neurons. For estimating the fraction of oracle performance for each model, we used the slope of a linear regression without offset fitted on the oracle correlation to each model’s test. Note that a network with shifter and modulator components could achieve a fraction of oracle performance greater than one, since the oracle is only conditioned on repeats of the same image and not other factors relating to brain state that the models could access. To avoid that confounding element, we froze the shifter and modulator input to the mean input across the training set and computed the fraction oracle based on the resulting network.

#### Selection of neurons for MEI generation

Following the training of the neural networks, we selected neurons to generate the MEIs based on the following criteria, in order:

- select neurons in the top 50^th^ percentile of oracle correlation to select cells with reliable responses to the image stimuli
- exclude neurons within 10 *µ*m of the edge of the scan field. This increases the chance of finding the same neuron across scans spanning multiple days.
- among these, select cells in the intersection of the top 30^th^ percentile of CNN model’s fraction correlation score *ρ*_CNN_ and top 30^th^ percentile of the LN model’s fraction correlation score *ρ*_LN_. This selects units that were predicted reasonably well by both CNN and LN models. In particular, it ensures that each neuron has a significant linear part that can be predicted by a quasi-linear model using the linear RF.
- order the remaining neurons from the largest to smallest in *ρ*_CNN_ *- ρ*_LN_. Iterate through the neurons, and place the visited neuron into a candidate set to be kept. At each iteration, remove any unvisited neuron that falls within 20 *µ*m distance of this unit, thereby excluding neurons that are too close to each other. This helps to reduce the chance of using same neuron more than once.

These selection criteria yielded total of 206 and 304 neurons from Mouse 1 and 2, respectively, ordered from the largest to smallest *ρ*_CNN_*-ρ*_LN_. Among these, the top 150 neurons were selected for MEI generation for each mouse.

#### Synthetic stimulus generation

To generate stimuli that optimally drive particular cells in primary visual cortex, we use methods similar to previous approaches in deep learning targeted at visualizing hidden features of deep networks (*16, 50–60*). For each model neuron, we construct an image that most strongly activates it (*58*), subject to a number of regularization constraints encouraging stable results. For each selected network architecture (CNN *vs* LN model for Mouse 1 or Mouse 2), we have four instances of the network trained to predict the neuronal population responses to images, *f* : ℝ^*w×h×c*^ *→*ℝ ^*n*^ but initialized with different random starting conditions. Given these models, we start with a Gaussian white noise input image *x*_0_ *∈* ℝ^*w×h×c*^ and optimize the image by iteratively perturbing the image with the gradient of the target neuron activity averaged over the 4 instances of the network with respect to the image ∇_*x*_*f.* Optimizing on four networks simultaneously helps to reduce high frequency noise that can vary significantly with the starting image and can obscure the generated image (*53*). We use two additional strategies to reduce the effect of the high frequency noise that is particularly prominent at the beginning of the gradient ascent procedure. Firstly, we apply Gaussian blur to the image after every gradient ascent step, starting with large value of standard deviation, and gradually decreasing over iterations (*61*). Secondly, we precondition the gradient (*53*) at each iteration by smoothing it by the lowpass filter 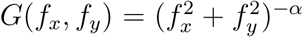 in the Fourier domain, which preferentially suppresses the higher frequency content of the gradient. We found *α* = 0.1 to be a good value by visual inspection of the resulting images.

When the above synthetic image generation technique is applied to a CNN model, the resulting image is a Most Exciting Image (MEI) for the target neuron. When the method is applied to a LN model, on the other hand, the resulting image is equivalent to a (highly regularized) linear receptive field (RF).

After the optimization we matched the mean luminance and contrast of the MEI and RF to ensure a fair comparison (111 mean pixel intensity and 16 RMS).

#### Synthetic stimulus presentation

In each of the closed-loop scan, we presented 150 MEIs and 150 RFs repeated 20 times each, exactly as generated but upscaled to the monitor resolution using 3^rd^ order spline interpolation. To detect potential misalignment in the screen position across multiple days, we presented 8 copies of the 150 MEIs shifted by *±*3.3º horizontally and vertically with no repetition. Hence a total of 7200 images were presented in the closed-loop scan. Just as in natural images, each image was presented for 500 ms followed by a blank screen with duration uniformly distributed between 300 ms and 500 ms.

#### Data analysis

The recorded responses of the matched neurons to the MEIs and RFs were normalized by diving each neuron’s response by the standard deviation across all presented MEIs and RFs, including the repeats and the shifts. We performed this normalization separately for each scan. The normalized responses of the matched neurons were then pooled across multiple MEI/RF presentation scans for each mouse (2 MEI/RF presentation scans for Mouse 1 and 3 MEI/RF scans for Mouse 2, all occurring on separate days), and averaged across repeats for each distinct image (Fig. 4A, Fig. S6).

To assess whether the MEIs generated for their target neurons tend to elicit higher responses than their corresponding linear RFs, we compared the average response across repeated presentations of the neuron’s MEI to the average response across repeated presentation of the neuron’s RFs. The overall difference in the average responses to MEIs *vs.* RFs pooled across all 300 neurons and both mice were assed using Wilcoxon Signed-Ranks test (*Z* = 5816.0, *p* = 7.6 *×* 10^-29^). For single neurons, the statistical significance in the difference in response to MEI vs. RF was assessed using Welch’s *t*-test with average d.o.f. of 34.5 across the pooled repeats (38–40 repetitions for Mouse 1 and 58–60 repetitions for Mouse 2 MEI/RF images).

### Supplemental Analysis

#### Difference in spatial frequency content between MEIs and RFs

We compared the spatial frequency content of MEIs and RFs by computing the average differences in the amplitude of the spatial frequency spectrum of the MEI and RF images,

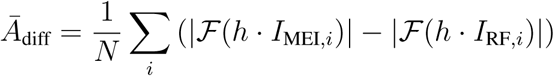

**Figure S2:**
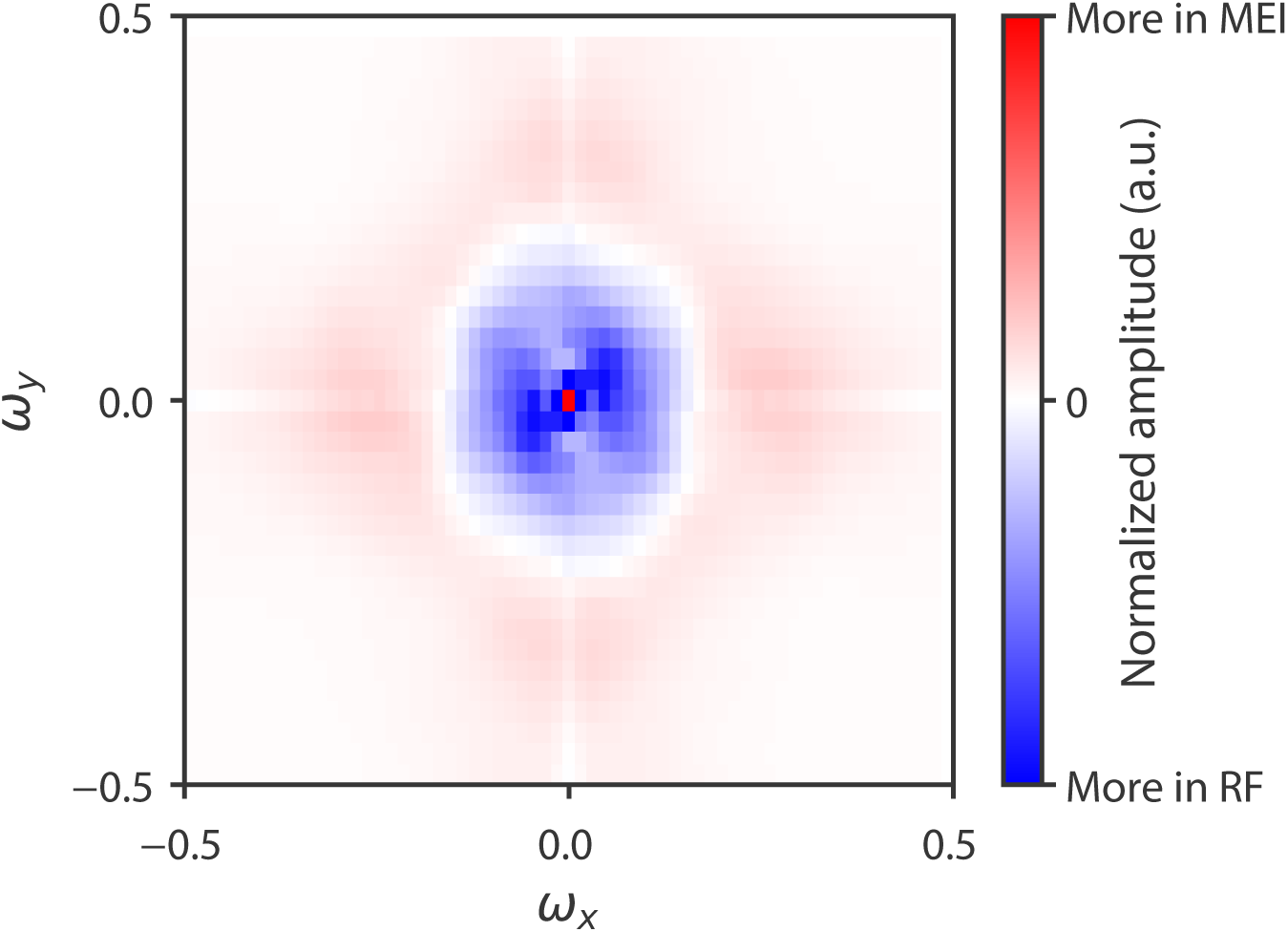
MEIs have higher spatial frequency content than RFs. The average difference in the amplitude of spatial frequency spectrum of MEIs and RFs. Positive value (red) indicates spatial frequency content that is, on average, stronger in the MEIs.

where *ℱ*(*·*) denotes 2D Fourier transform, *h* is a Hamming window and *N* is the total number of MEI/RF image pairs.

#### Non-trivial nonlinearity of the CNN

A linear-nonlinear (LN) model has the form *y* = *g*(**w**^*T*^**x** + *b*) where *y* is the neuronal response prediction, **x** is the input image, **w** is a weight vector of the same dimensions, and *b* is an offset. *g* is a static non-linearity which is usually chosen as part of the model design. In our case, we chose *g*(*z*) = ELU(*z*) + 1.

It is possible that the CNN model differs from the LN model only in trivial ways by effectively learning an LN model but achieving higher prediction scores because of more freedom in fitting *g* or because of an in-built architectural bias that makes learning the **w** easier.

To demonstrate that the CNN model deviates non-trivially from the LN model, we computed the gradient of both models with respect to the input image on the entire image dataset and computed the largest 10 eigenvalues of the covariance matrix of these gradients. For the LN model all gradients are proportional to **w**. Thus there must be exactly one eigenvalue greater than zero. If the CNN model just behaves like a linear-nonlinear model, the spectrum should look the same. However, if it is nonlinear in a non-trivial way, the gradients should differ and the spectrum should have more non-zero eigenvalues.

We find the latter to be the case (Fig. S3), indicating that the CNN model performs better because it can match interesting nonlinearities of cortical neurons that cannot be captured by the linear model.

**Figure S3:**
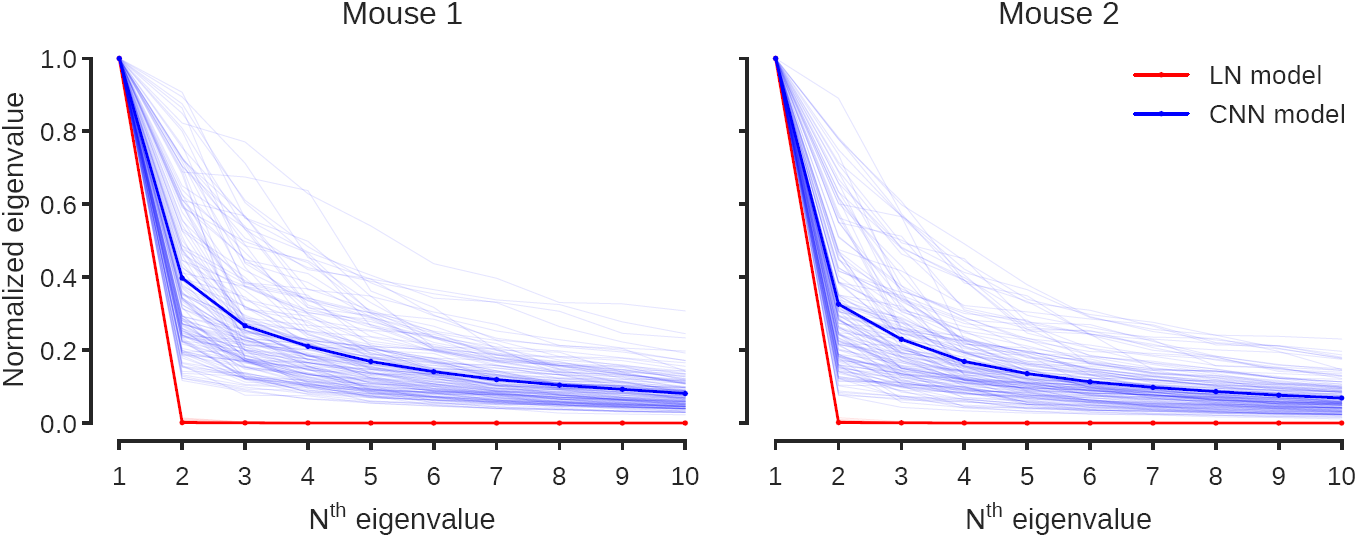
The CNN models are nonlinear in non-trivial ways. The two plots show the first ten eigenvalues of the covariance matrix of the gradients of the CNN model and the linear-nonlinear model on the entire image set. Different spectra correspond to different neurons. Each spectrum was normalized to the largest eigenvalue. As expected the linear-nonlinear model has a one-dimensional gradient spectrum. However, the CNN model has several eigenvalues greater than zero, demonstrating that it is nonlinear in a non-trivial way.

## Supplemental Figures

**Figure S4:**
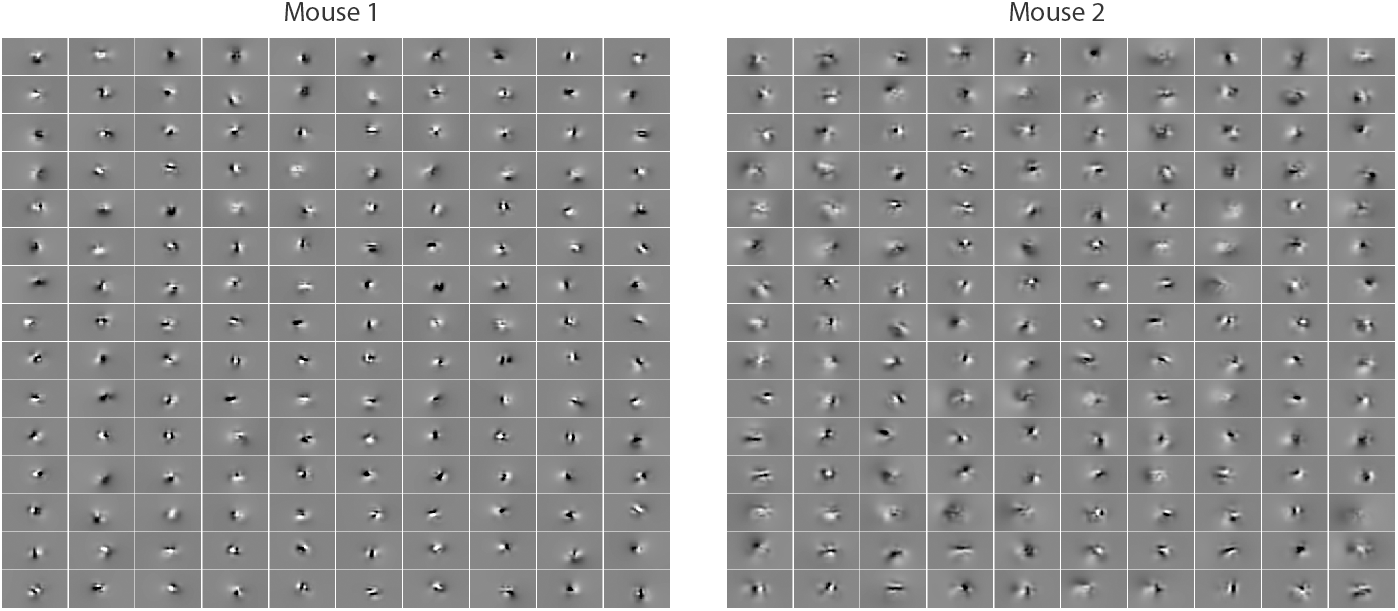
All linear receptive fields for all 300 neurons used in the close loop experiment for both mice.

**Figure S5:**
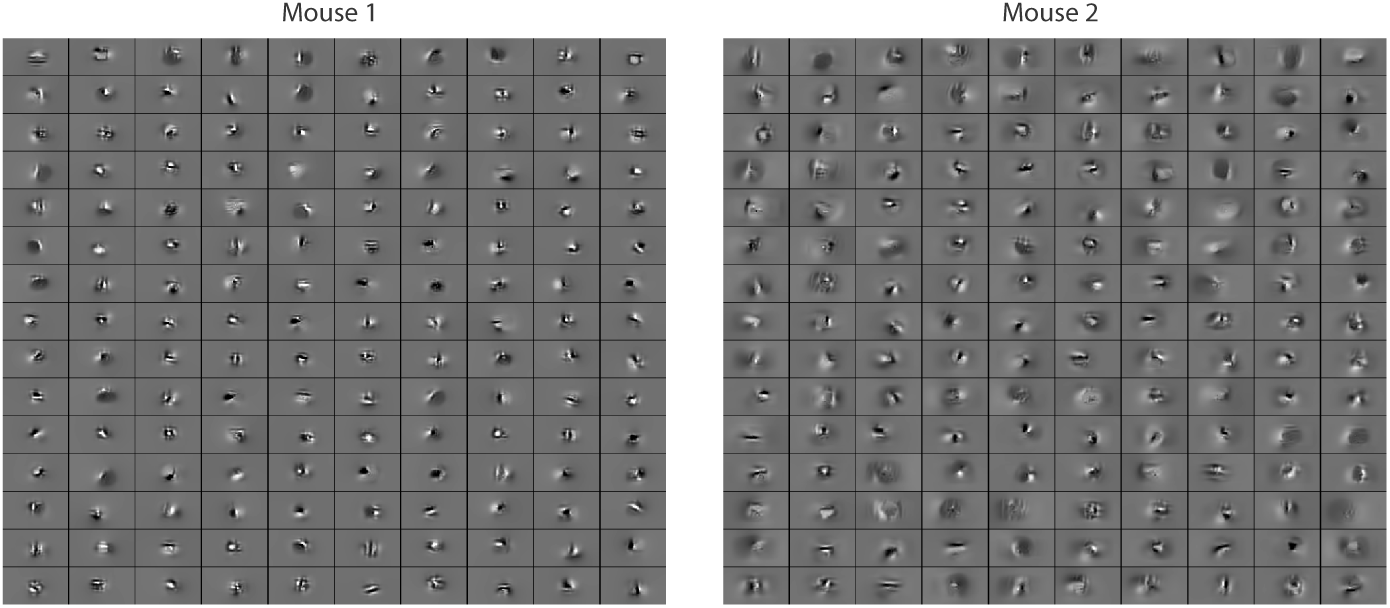
All MEIs for all 300 neurons used in the close loop experiment for both mice.

**Figure S6:**
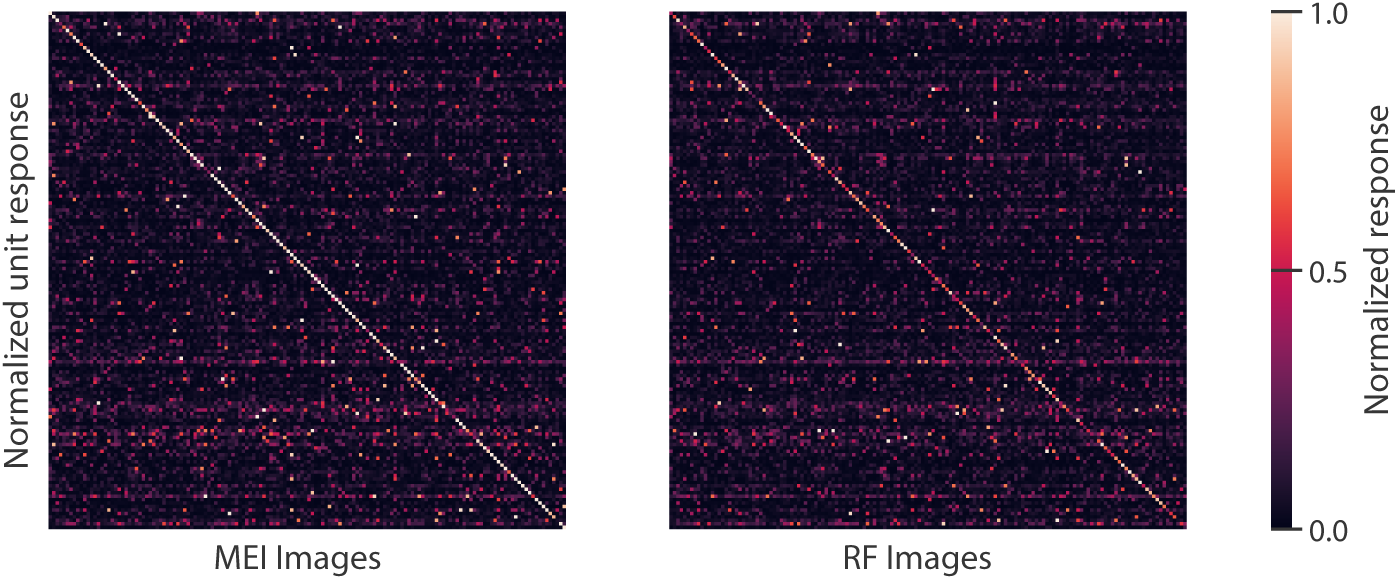
Equivalent of Fig. 4A for Mouse 1.

